# Fixation-evoked potentials reveal neural signatures of hierarchical value-integration during decision-making

**DOI:** 10.64898/2026.07.15.738682

**Authors:** Jordan Deakin, Romy Frömer, Sebastian Gluth

**Author notes:** Corresponding authors: Sebastian Gluth; Jordan Deakin, University of Hamburg, Von-Melle-Park 11, 20146 Hamburg, Germany.

## Abstract

Many everyday decisions require evaluating and weighting information across attributes, yet it remains unclear whether the brain compares attributes directly or first integrates them into option-level representations. Combining EEG and eye-tracking during naturalistic multi-attribute choice, we derived fixation-level predictions from a hierarchical Bayesian model and tested them against fixation-evoked neural activity. Behavioral and eye-tracking data confirmed that search and choice dynamics were well predicted by the model and thus more consistent with a value-integration as compared to a pure attribute-comparison strategy. Crucially, we found distinct neural signatures mirroring the model’s processing hierarchy. A fronto-central positivity reflected attribute-level belief updates, whereas option-level updates were tracked by a centro-parietal positivity. Critically, only the option-level signal scaled with attribute importance and predicted subsequent choice. Furthermore, lateralized preparatory motor activity tracked the evolving preference based on this integrated value difference. These findings provide neural evidence for a hierarchical, integrative architecture of multi-attribute choice and establish a general framework for studying decision computations under naturalistic viewing conditions.

---

A defining feature of human cognition is to excel in making decisions with high complexity that unfold through the extended sampling, weighting and integration of information. Therefore, a central challenge for theories of decision-making is to explain how information is sampled and combined to form preferences^1,2^. Converging evidence from eye-tracking studies investigating both value-based and perceptual decisions^3–6^ has shown that attention and preference are tightly coupled. Decision-makers more often fixate items deemed attractive^7,8^ and are also more likely to choose an item that they looked at longer^5,6,9,10^. This bidirectional relationship between (overt) attention and choice suggests that preferences emerge dynamically as information is sampled over time.

Accompanying these empirical findings, computational models have helped formalizing the interaction between attention and choice in simple perceptual and value-based choices^3,6^. Many decisions, however, require more extended deliberation, during which information is actively sampled across multiple sources before being integrated over time. Arguably the most prominent example of such complex decision-making is multi-attribute choice, which has been formalized in many extant computational models^6,7,10–15^. These models range from variations of the attentional Drift Diffusion Model (aDDM), which treat attention as a bias on evidence accumulation in favor of attended attributes and options^6,10,16^, to theories that explicitly model attentional allocation, for example by assuming that attributes are sampled in accordance with optimal search policies that maximize expected utility under sampling costs^17^. Critically, these models differ in their assumptions of how values are computed and compared across options to make a choice^18,19^. Some accounts propose that attributes are integrated into single value representations for each option, which are compared to determine the choice^6,10,20,21^. In contrast, other accounts propose that isolated option-level representations may not exist at all, with choices arising directly from attribute-level comparisons^13^ or from applying heuristic strategies^22^.

Attempts to elucidate this question on the neural level have been sparse, with most neuroscientific work disproportionately relying on simple choice tasks^23–27^. Moreover, research extending this approach to multi-attribute choice has been limited largely to non-human primates^28–31^ or human fMRI studies^30,32–35^, aimed at identifying brain regions encoding either the integrated value of options^30^ or comparisons between their constituent attributes^31,35^. While EEG offers substantially greater temporal resolution, it has traditionally been difficult to apply to tasks involving active information search. Under naturalistic conditions, decision-makers actively sample information through eye movements, yet these eye movements introduce substantial artifacts into EEG recordings^36,37^ and, perhaps more importantly, induce temporal variability in processes of interest and their associated neural activity across trials. To circumvent these issues, previous EEG studies have constrained information search by artificially enforcing central fixation and short response deadlines^24,25,38^. Although these restrictions facilitate the identification of choice-related neural activity, they constrain the decision process in ways that depart from how such complex choices are typically made, where information is actively sampled through shifts of gaze and attention. Consequently, the neural mechanisms underlying multi-attribute choice remain poorly understood. Bridging this gap requires combining computational models that generate quantitative predictions about latent decision processes with methodologies capable of measuring neural activity during active information search.

Here we address this gap in two ways. On the computational level, we use our recently developed Multi-Attribute Search and Choice theory (MASC)^21^, a hierarchical Bayesian belief-updating model which predicts how information is sampled and integrated in complex decision problems. MASC assumes that decision-makers maintain probabilistic beliefs about the value of attributes and options, formalized as hierarchical belief distributions. Attention guides the sampling of attribute information, which is then weighted and integrated into option-level beliefs that are compared to determine a choice. In this way, MASC adopts and ‘integrate-then-compare’ account of multi-attribute choice, distinguishing it from purely comparison-based theories in which choice emerges directly from attribute-level comparisons^18,19^ as well as from theories proposing (attention-weighted) parallel processing of all available information^10,39^. Central to the model is its myopic search rule, which assumes that observers plan one step ahead, sampling the attribute that maximizes the probability of choosing the associated option at the next step^12,40^. Despite its simplicity, this search rule gives rise to adaptive search behavior in which attention is guided by a combination of value, uncertainty and attribute importance. In doing so, MASC accounts for a range of empirical phenomena in both simple and complex decisions, including the bidirectional relationship between attention and choice^3,5,6,8^ and the distribution of attention to attributes over time^7,10,41,42^.

On the neural level, we leverage MASC to derive and test predictions for neural activity by combining EEG and eye-tracking in a pre-registered study. Combining EEG and free-viewing paradigms presents substantial challenges (e.g., the correction of ocular artefacts, temporal variability in process-relevant neural activity and the temporal overlap between fixation-evoked responses). To overcome these challenges and link the momentary sampling of information to corresponding neural processes observed during the decision process, we harnessed recent advancements in EEG and eye-tracking coregistration and analysis techniques enabling the isolation of neural activity attributable to individual fixations (fixation-evoked potentials, FEPs)^43–47^. We derived fixation-level predictions from MASC and tested whether these predictions could explain variability in the corresponding FEPs. Given MASC’s distinction between attribute-level and option-level belief updates, this allows us to test whether the proposed Bayesian belief-updating dynamics are reflected in the brain and critically, whether attributes are integrated into option-level values that subsequently form the basis of comparison. In accordance with our model, we predicted to identify transient belief-updating signals at both levels of the choice hierarchy (i.e., attributes and options), as well as more sustained signals of emerging preference based on the integrated value difference between options.

In addition to replicating our previous work that MASC accounts for a multitude of attention-choice interactions^21^, we show that MASC-derived belief updates are indeed predictive of fixation-evoked neural activity, with dissociable neural signatures for attribute and integrated option-level belief updates. Importantly, only the strength of the option-level belief updating signal depends on the importance of the sampled attribute and is predictive of the subsequent choice. Together, our results provide converging behavioral and neural evidence that multi-attribute decisions involve sequential, fixation-driven belief updating consistent with Bayesian inference, whereby sampled attributes are weighted and integrated into option-level representations forming the basis of comparison. More broadly, our findings establish a link between computational models of information search and the neural mechanisms that support complex decision-making and showcase how to investigate the neural basis of attention and choice dynamics under naturalistic free-viewing conditions.

## Results

### MASC captures the interplay of fixation and choice dynamics

We recorded EEG and eye-movement data while participants (N = 57) made self-paced choices between pairs of smartphone options. Each smartphone was characterized by star ratings (0-5) for three attributes: screen size, battery capacity and storage space. Choice sets comprised an equal number of easy, medium and hard trials tailored to each participant based on item ratings obtained immediately before the choice task (Fig. 2a). In the main task, participants indicated their preferred option via button presses, using their left and right index fingers for the left and right options, respectively. To derive the model-generated estimates used to predict fixation-evoked activity, we fit the MASC model to the behavioral data (see Methods for details) and verified that the model captured the observed search patterns and choices. In brief, participants’ behavior replicated previous findings^21^ and was well accounted for by MASC (Fig. 3).

**Fig. 1.**
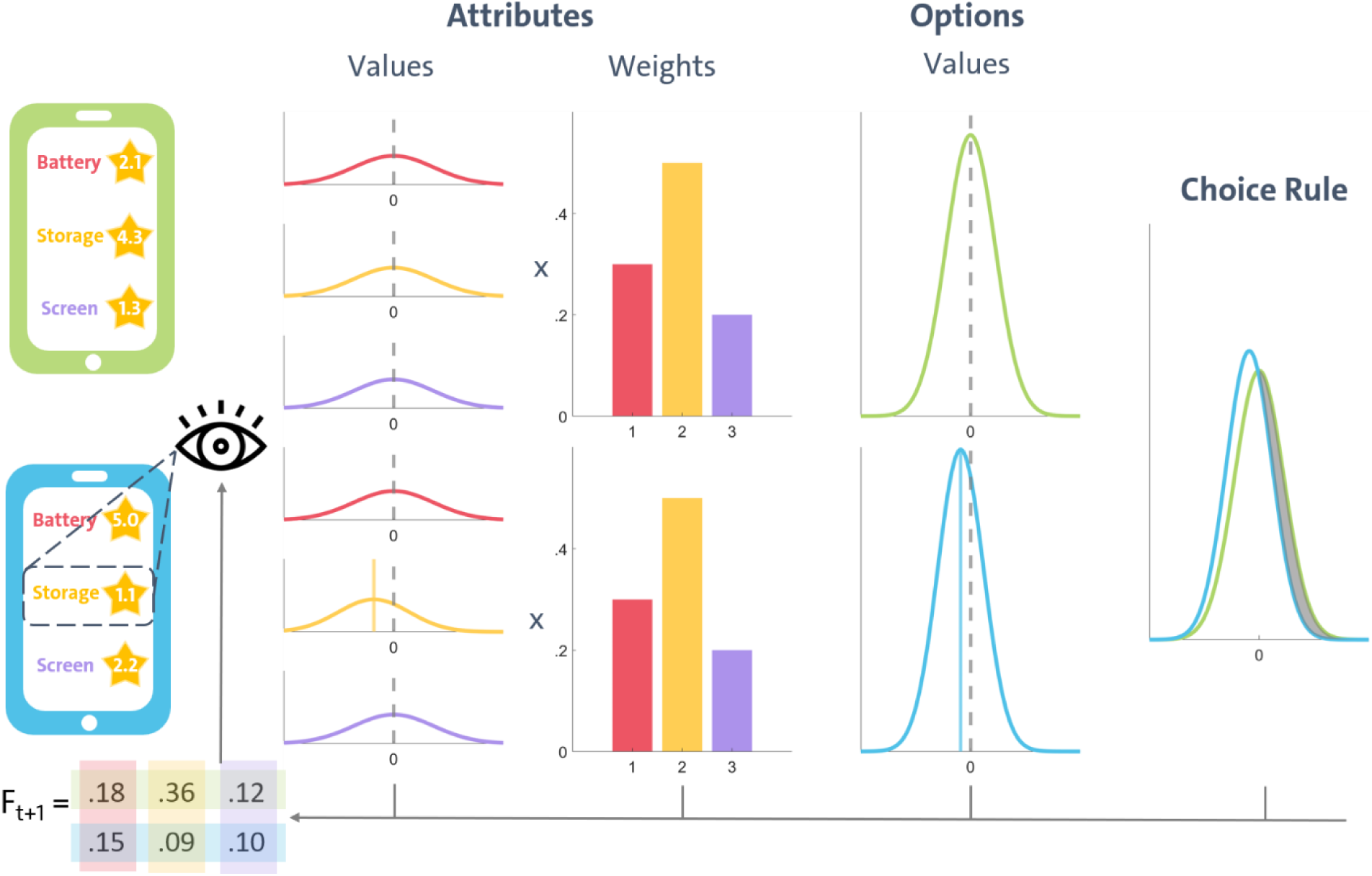
Schematic diagram of Multi-Attribute Search and Choice (MASC). Beliefs about the values of attributes and options are formalized as hierarchical belief distributions. Guided by the myopic search rule, the decision-maker selects an item on which to fixate. Thus, the high-weighted ‘storage’ attribute is likely sampled first (as shown here), and a sample is obtained which suggests a comparatively low attribute value. The corresponding attribute and option distributions are updated accordingly. After one fixation, the small separation between option posteriors does not constitute sufficient evidence for a choice, so a new fixation is generated. The model predicts that the observer will likely switch to the green option (p = .18 + .36 + .12 = .66) which currently has both higher value and higher uncertainty. Sampling its high-weighted storage attribute is most likely (p = .36). Taken together, it emerges from MASC’s myopic search rule that weights, values, and uncertainty drive the information sampling process.

**Fig. 2.**
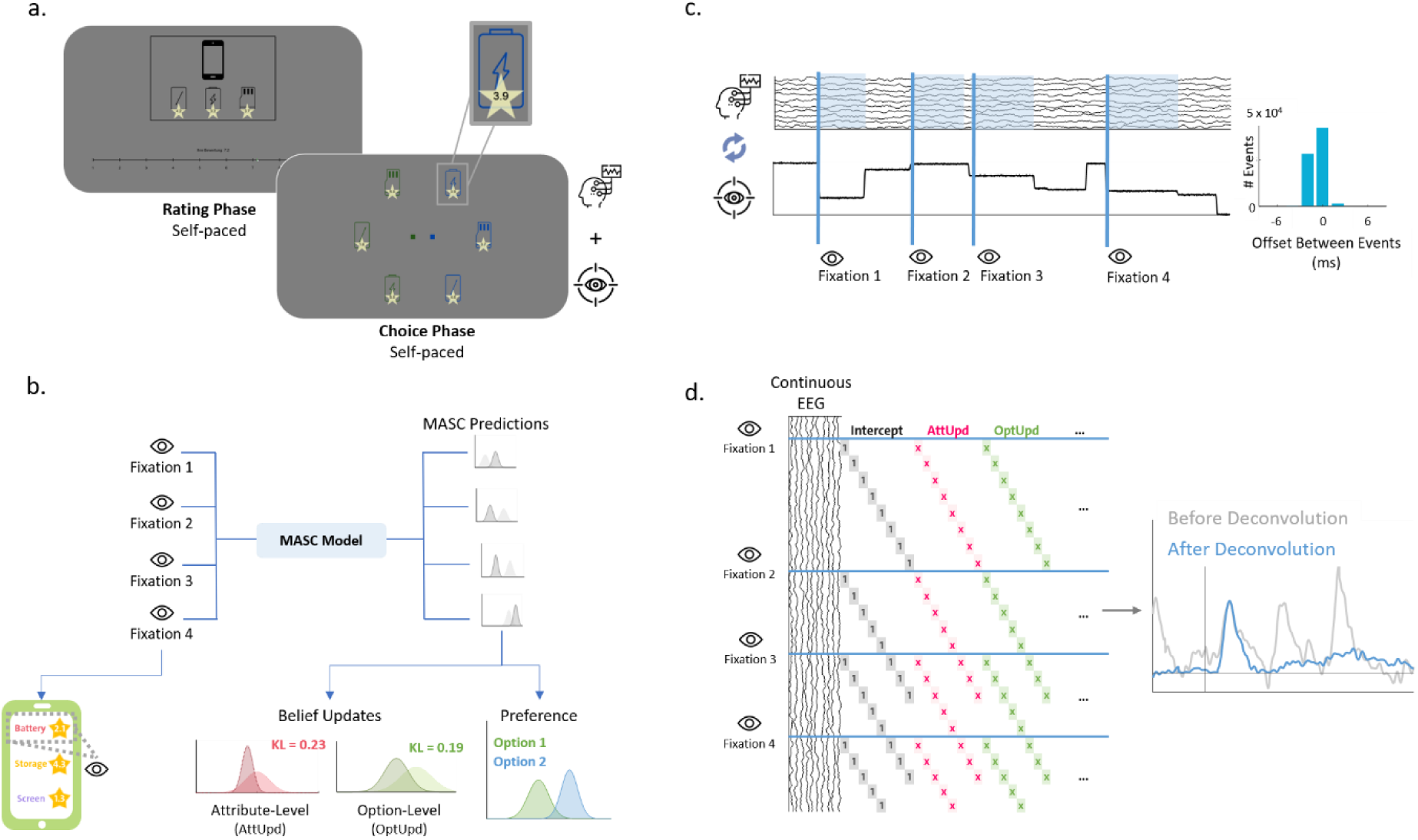
Experimental design and data analysis pipeline. **a.** Upper panel: Trial structure for the rating task. Participants rated 45 smartphones, each characterized by star ratings (0-5) for three attributes. Icons for each attribute were first presented without star ratings (1500 ms) and then together with the star ratings and rating scale (1-10). Lower panel: Trial structure for the choice task, in which participants chose between pairs of smartphones. The six attribute icons (three per option) were first presented alone (for 1500 ms) before the star ratings appeared. Options were separated by color (green vs. blue) and position (left vs. right). Participants were free to move their eyes and choose their preferred option at any time. **b.** Generating MASC predictions: The model was fitted to the choice and eye-tracking data and simulated with the best-fitting parameters for each participant. For each fixation, the belief distributions were updated in accordance with the fixated attribute. Belief updates at the attribute (option) level were quantified by the KL divergence between the attribute (option)-level posterior pre vs. post fixation. Preference was quantified as the Jeffrey’s distance between posterior option distributions after the fixation. **c.** Synchronizing EEG and eye-tracking. Left panel: The two data streams were synchronized based on shared triggers sent to each system. Fixation onsets from the eye-tracker were used to mark corresponding events in the EEG signal. Right Panel: Across all participants and events, the average synchronization error was < 2 ms. **d.** Linear deconvolution allows modeling each fixation as a combination of MASC-generated predictors (left panel), while also removing temporally overlapping activity from neighboring fixations (right panel).

**Fig. 3.**
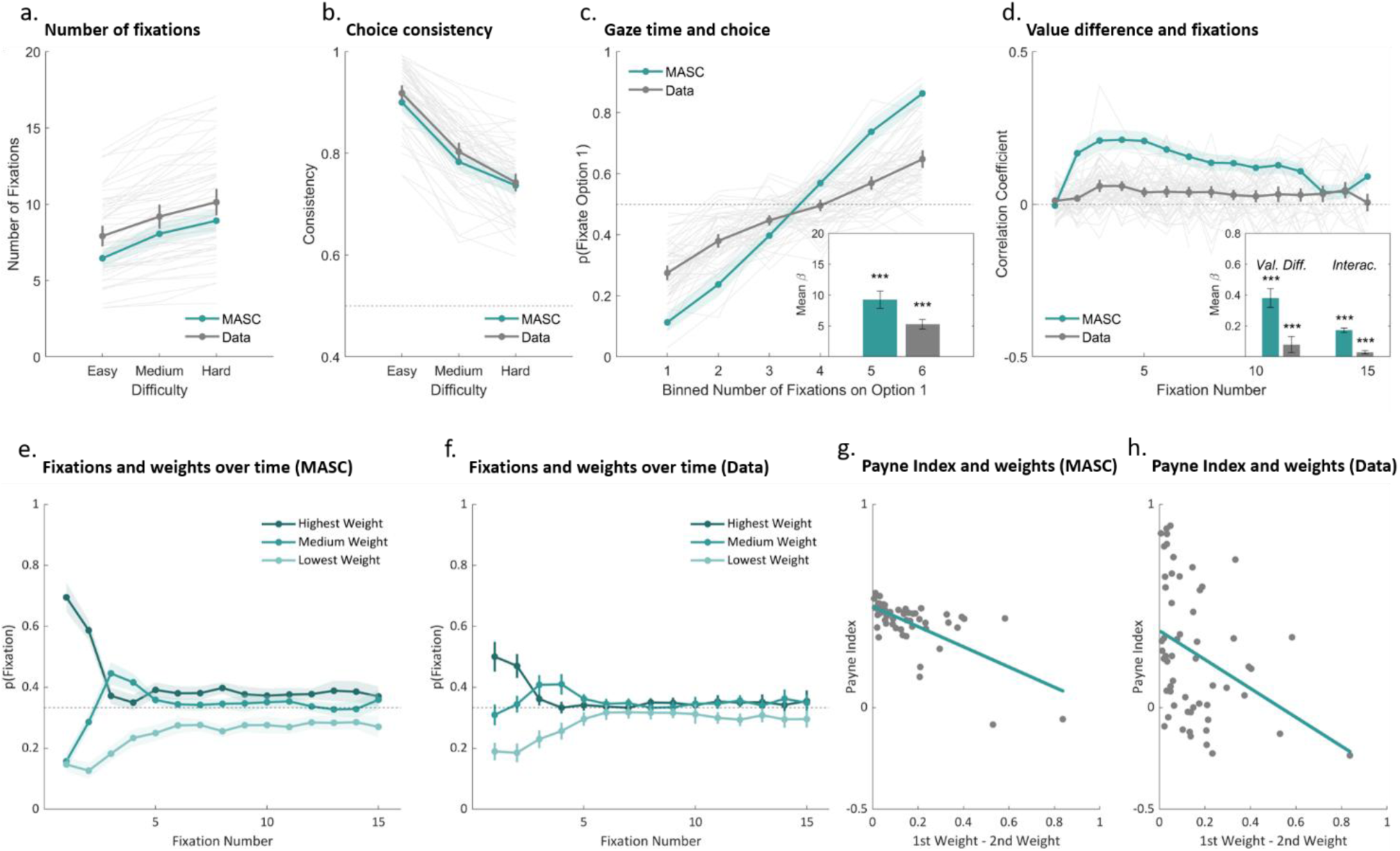
Behavioral and eye-tracking data with corresponding MASC predictions. **a.** Difficult trials require more samples to tease apart options, leading to an increase in the number of fixations with increasing difficulty. **b.** Efficient searches are non-exhaustive, leading to lower choice consistency in difficult trials. **c.** Sampling positive information from a choice option promotes further inspection of it and increases its choice probability, leading to a positive association between attention and choice. **d.** Attending to high-value items increases the probability of making a choice in the next step. As such, higher-value options are attended to more often (Val. Diff. = value difference), and this tendency increases over time (Interac. = interaction). **e.-f.** Important attributes are attended early on, as they provide more decisive evidence. Later, attention shifts to other attributes as uncertainty reduction and value take precedence in guiding attention. **g.-h.** Participants who assign disproportionately high weight to one attribute tend to make more attribute-wise transitions, as reflected in a lower Payne Index. Shaded areas and error bars represent 95% confidence intervals; ***p < .001.

First, MASC predicts an increase of the number of fixations and a decrease of choice consistency with increasing difficulty (i.e., decreasing option-value difference). Within MASC’s Bayesian framework, increased difficulty corresponds to greater overlap between the two posterior option distributions, in which case more samples are required to disambiguate the options. Yet, this sampling increase is rarely exhaustive, so that consistency is still predicted to decline for more difficult trials. In line with the model’s predictions, participants made fewer fixations in easy compared to medium difficulty trials (*t*(56) = -10.082, *p* < .001, Cohen’s *d* = -1.335) and in medium compared to hard trials (*t*(56) = -9.450, *p* < .001, *d* = -1.252, Fig. 3a). Moreover, participants chose the higher-value option more frequently in easy compared to medium difficulty trials (*t*(56) = 18.290, *p* < .001, d = 2.423) and in medium compared to hard trials (*t*(56) = 7.840, *p* < .001, d = 1.038, Fig. 3b).

Previous work has shown that options receiving more attention are more likely to be chosen^4,6^. MASC reproduces this effect, which emerges naturally from its search rule, because early positive samples promote further inspection of the same alternative (in an effort to obtain decisive evidence), thereby increasing its choice probability. To test this prediction, we regressed the probability of choosing Option 1 onto the proportion of fixations allocated to Option 1 while controlling for value difference in a logistic regression. In accordance with MASC, this analysis revealed a positive association between attention and choice probability (*t*(56) = 13.730, *p* < .001, *d* = 1.819, Fig. 3c). The tendency to revisit promising options demonstrates that attention is guided by value in MASC, in line with studies demonstrating a positive association of value and fixation that strengthens over time^7,42^. To test for this effect in the data, we regressed the probability of fixating Option 1 onto the value difference between the two options, the fixation number and their interaction. The positive main effect of value difference showed that participants indeed sampled higher value options more often (*t*(56) = 10.755, *p* < .001, *d* = 1.425), while the significant interaction confirmed that this bias increased over time (*t*(56) = 5.146, *p* < .001, *d* = 0.682, Fig. 3d).

As well as prioritizing high values, MASC also predicts an intricate dynamic dependency of attention on the importance of information (i.e., attribute weights). The decision-maker is expected to first focus on the most important information but then shift attention to less important information when the former provides insufficient evidence (Fig. 3e). The empirical patterns in our experiment are strikingly similar to the predictions of MASC (Fig. 3f). To test this formally, we regressed the probability of fixating an attribute as a function of its weight and fixation number. The intercept coefficient was significantly higher than chance level, showing that participants fixated more often on the highest weighted attribute (*t*(56) = 6.380, *p* < .001, *d* = 0.845). Likewise, the effect of fixation number was negative (*t*(56) = -4.197, *p* < .001, *d* = -0.556), showing that this tendency decreased over time.

Finally, we tested whether MASC captures individual differences in search patterns as a function of the dispersion of attribute weights. Previous work has shown that participants with more extreme attribute weights tend to exhibit more attribute-wise than option-wise search patterns^10^. MASC predicts this effect because comparing a dominant attribute across options should provide the most decisive evidence (Fig. 3g). To test this prediction in the data, we examined the correlation between attribute dispersion (defined as the difference between the two highest-weighted attributes) and the Payne Index^48^, which summarizes search patterns by calculating the ratio of option-wise to attribute-wise transitions. In line with MASC’s prediction, this correlation was significantly negative (*r*(55) = -0.349, *p* = .004, Fig. 3h). Notably, the model also correctly predicts that option-wise transitions were more common than attribute-wise transitions, as indexed by the majority of participants exhibiting a positive Payne Index.

Taken together, we replicated the series of behavioral and eye-tracking results reported in our previous work^26^, suggesting that multi-attribute decisions are governed by efficient information-search and preference-formation dynamics as predicted by our theory. In particular, the modulation of search patterns by attribute-weight dispersion, together with the predominantly option-wise nature of search overall, is consistent with MASC’s assumption that attribute samples are integrated to form and compare option values.

### Analyzing fixation-evoked EEG potentials

Having established that our computational framework accounts for the empirical search and choice patterns, we next examined the neural dynamics predicted by MASC. More specifically, our theory assumes that decision-makers sequentially fixate attributes to sample their values (i.), update attribute-levels beliefs based on this information (ii.) weight and integrate these beliefs into option-level representations (iii.) and compare these integrated representations to determine a preference for the currently favored option (iv.). We therefore predicted to identify latent belief-updating signals at both levels of the choice hierarchy (i.e., attributes and integrated options) that are not only separable from each other but also distinct from signals reflecting the fixated attribute values as well as the emerging preference. Thereto, we defined the following linear deconvolution model^44,49,50^ to predict fixation-evoked EEG activity

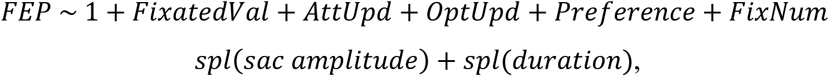

where *FixatedVal* refers to the value of the fixated attribute, *AttUpd* and *OptUpd* are MASC-generated predictors (based on the Kullback-Leibler divergence) quantifying the magnitude of belief updates at the attribute and option levels, respectively, and *Preference* quantifies the strength and direction of evidence for one option over the other (see Methods for details). *FixNum* controls for the ordinal fixation number within a trial and the final two predictors (sac_amplitude and duration) are spline covariates used to control for the non-linear effects of saccade amplitude and fixation duration on the FEPs^47,51–54^. After fitting the deconvolution model to each participant separately, the resulting beta weights were subject to threshold-free cluster permutation tests^55,56^ to identify timepoints and electrodes where these predictors explained the most variance. Strikingly, these tests revealed significant main effects of all predictors, with distinct topologies and temporal characteristics for each (Fig. 4).

**Fig. 4.**
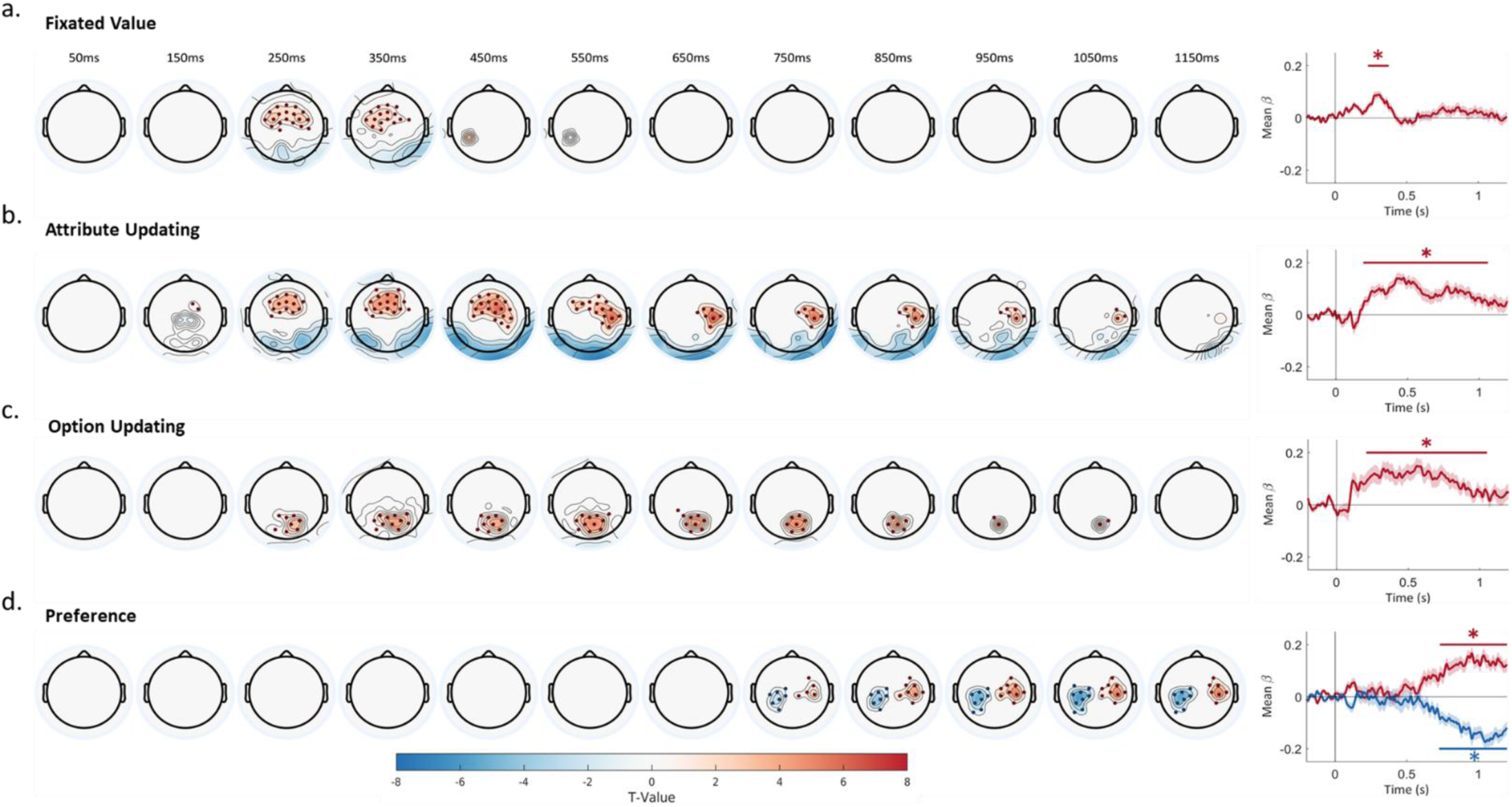
MASC-generated predictors for FEPs reveal distinct spatiotemporal profiles. Topographical maps showing the t-values of significant clusters associated with the predictors of fixated value (**a.**), attribute-level updating (**b.**), option-level updating (**c.**), and the preference of the currently favored option (**d.**). Statistical maps are obtained using threshold-free permutation testing (E = 0.5, H = 2) and averaged across a 100 ms sliding window (left). The right panels show the beta coefficients averaged across channels within the respective cluster. Lines indicate the temporal extent of significant clusters.

### A transient and non-linear representation of fixated value

As a first processing step at each fixation, the decision-maker is expected to process the sampled attribute’s value. This predictor was associated with a transient effect spanning 230-374 ms (peak latency = 306 ms, p < .001) in a cluster of fronto-central electrodes, indicating more positive amplitudes for higher value attributes (Fig. 4a). To explore the effect in more detail, we reconstructed the individual FEPs using the beta weights from the deconvolution model and averaged them according to the rounded value of the fixated attribute (Fig. 5a). Notably, this analysis suggests the fixated-value effect to be primarily driven by fixations to the highest value attributes (star ratings 4-5), with little difference between average and low values (Fig. 5b). High-value attributes are not only relevant for making a choice based on momentary evidence but also play a key role in guiding future attention by facilitating the identification of promising options that merit further inspection. In this sense, high-value attributes may signal progress towards the goal of reaching a decision and thereby be subject to additional processing. Aligned with this interpretation, an analysis of fixation duration revealed a small but significant effect of value, with longer durations for higher values (Fig. 5b inset). Like the effect on the FEPs, this was driven by slightly longer fixations to the highest value attributes, with little difference between low and average values (see Supplementary Table 1-2 for statistical analyses).

**Fig. 5.**
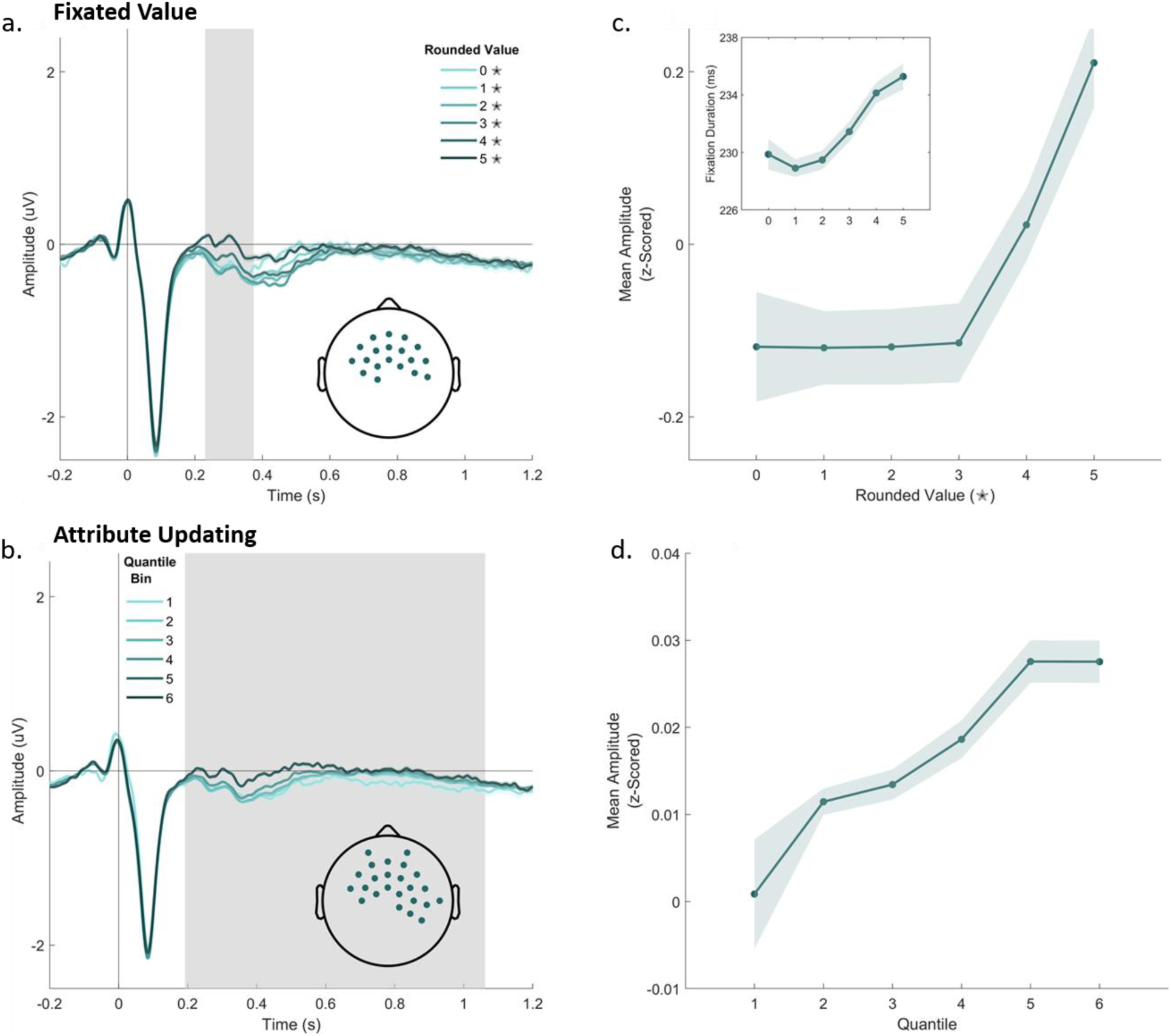
Reconstructed FEPs and average amplitude for the fixated value and attribute updating predictors. **a.** Reconstructed FEPs averaged across the electrodes in the FixatedVal cluster. Lines correspond to the average FEPs for the rounded value (star rating) of the fixated attribute, and the grey shaded area shows the time window associated with the cluster. **b.** Lines correspond to the average FEPs for the quantile (.1,.3,.5,.7,.9) binned level of the AttUpd predictor. **c,d.** Average amplitude of each average FEP within the time window associated with the cluster. For FixatedVal, the inset shows the effect of FixatedVal on fixation duration. Error bars represent the within-subject standard errors computed with the Cousineau-Morey correction.

### A fronto-central signal of sustained and linear updating of attributes

A core prediction of MASC is that sampling information leads to belief updates about the corresponding attribute values, shifting both the mean and precision of their belief distributions. Similar to the fixated-value signal, the magnitude of attribute-level updating was also associated with a positive, fronto-central cluster of electrodes (Fig. 4b). Yet, the effect was much more sustained (range: 192 ms – 1062 ms, peak time = 590 ms, p < .001). In contrast to the non-linear encoding of fixated value (Fig. 5b), the amplitude of this FEP scaled linearly with the magnitude of attribute updating (Fig. 5d), suggesting an accurate neural representation across both smaller and larger belief updates. The temporal and topographical characteristics of this effect overlap with prior work showing that larger belief updates are associated with stronger fronto-central activity that scales with the degree to which internal models of the environment are updated^57–60^. Thus, it appears that the effects associated with fixated value and attribute-updating reflect two distinct but interlinked processes: initial appraisal of the fixated value triggering a corresponding attribute-level belief update.

### A hierarchical distinction between attribute and option-level updates

Once an attribute-level belief is updated, MASC assumes the next step in the hierarchy is to weight attribute-level beliefs to form beliefs about the option’s overall value. Notably, updating at the option-level is qualitatively different from attribute-level updating because it requires integration across multiple belief states. Consistent with this computational distinction, the effects for option updating were topographically distinct from those of attribute-level updates, with activity in a positive cluster of parietal electrodes being uniquely predicted by the option-updating predictor (range: 208-1050 ms, peak latency = 574 ms, *p* < .001). Centro-parietal activity has previously been linked to the accumulation of evidence in decision-making^61,62^. However, the present finding that this signal reflects *within-sample* integration of attribute information into option-level updates aligns most closely with a recent study linking the centro-parietal positivity (CPP) to similar sample-level belief updates in an expanded judgment task^63^.

### Only option-updating signals are dynamically shaped by attribute weights

A core feature of MASC is its ability to predict how the three factors of attribute weighting, uncertainty, and value dynamically guide attention during the emerging decision. Critically, the theory thus provides a precise account of the initial prioritization of the most important attribute, which then decays at later fixations (Fig. 3e-f). With respect to neural belief-updating signals, MASC predicts that attribute importance should modulate the magnitude of option-belief updates, since it assumes that option-level beliefs are a weighted combination of attribute-level beliefs. Taking both the attentional dynamics and the integration rule into account, we therefore expected the option-level updating signal to be more sensitive to attribute weights during early compared to late fixations. To test this prediction, we averaged the reconstructed FEPs across fixation number and the weight of the fixated attribute. Since the effect of attribute weight diminishes rather early, we restricted this analysis to the first six fixations of each trial. We took the time points and channels associated with option-level updating effects (Fig. 4c) and calculated the mean amplitude of the FEPs within the time windows of each cluster. These amplitudes were then subject to a 3 (Attribute Weight: highest, medium, lowest) x 6 (Fixation Number: first to sixth) repeated-measures ANOVA to test for both main effects and the interaction of the two factors. Notably, attribute weight was not a predictor in the deconvolution model and thus did not directly influence the estimates of the FEPs.

The ANOVA revealed all effects to be significant, and the mean amplitudes (Fig. 6b) closely resembled attentional dynamics (Fig. 3f). There was a main effect of Fixation Number (*F*(5,280) = 7.712, *p* < .001, η_p_^2^ = .121), with mean amplitude generally decreasing from the first to the sixth fixation, mainly driven by larger amplitudes for the first fixation. Crucially, there was also a main effect of Attribute Weight (*F*(2,112) = 7.935, *p* < .001, η_p_^2^= .124), such that option-update effects were larger for more important attributes. In line with the decaying prioritization of the most important attribute in both MASC and the eye-tracking data, we observed a similar development of the FEP amplitudes, resulting in a significant interaction between Attribute Weight and Fixation Number (*F*(10,560) = 2.383, *p* = .009, η_p_^2^ = .041). Simple effects analyses revealed that the influence of Attribute Weight was significant only at the first, second and fifth fixations, with a dynamic shift of the highest mean amplitude from the most important (at fixations one and two) to the second-most important attribute (at fixation five).

**Fig. 6.**
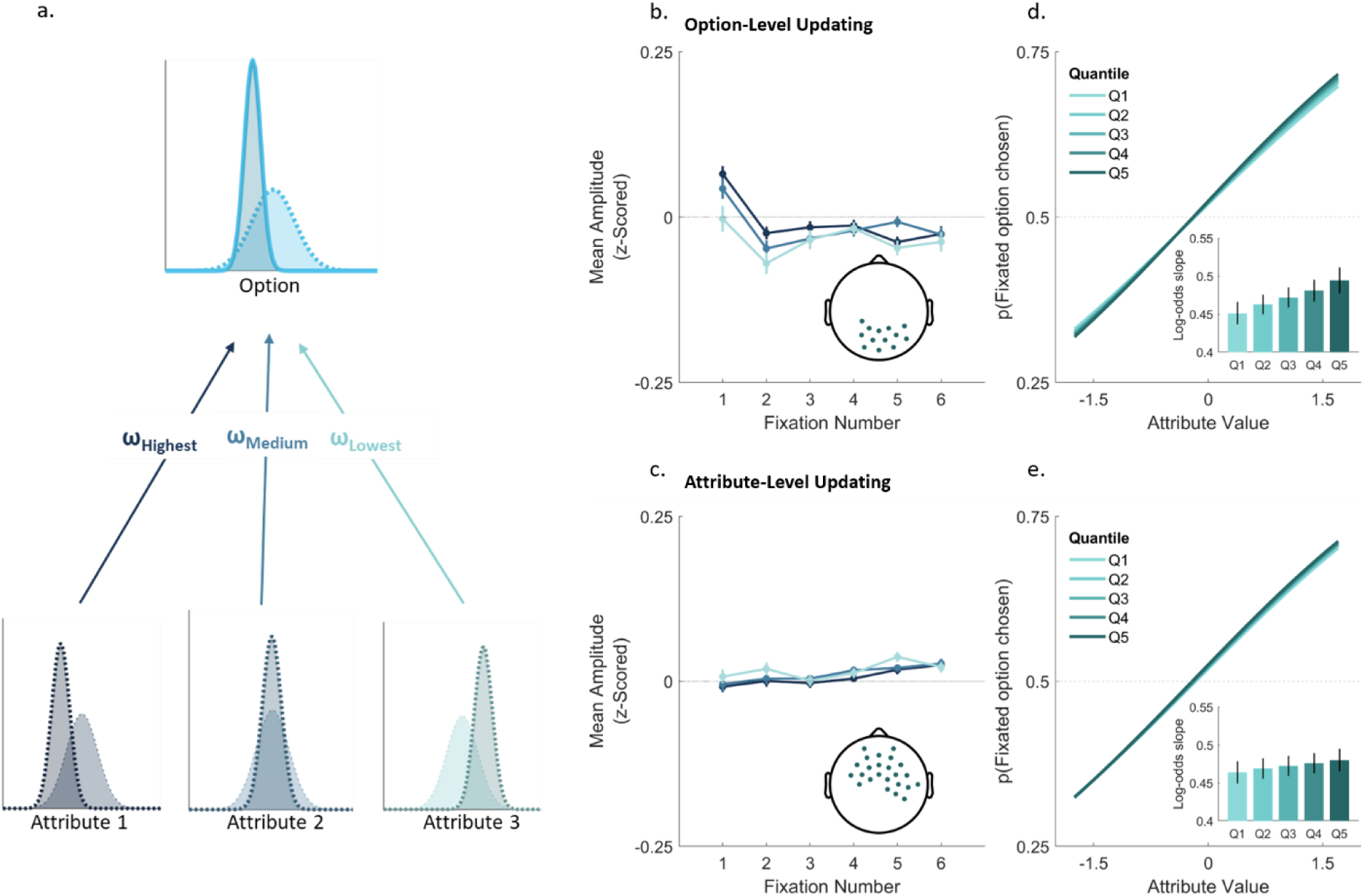
Effects of attribute weights on attribute-level and option-level updates. **a.** Hierarchical belief updating in MASC. MASC assumes attribute-level beliefs are weighted by importance and integrated to form option-level beliefs. In this example, the three attributes are believed to have comparatively low, average and high values, respectively. However, the highly weighted low-value attribute dominates, skewing the option distribution leftwards towards lower values. Thus, attribute weighting affects beliefs only at the option level. **b-c.** Mean z-scored amplitudes within the cluster time window for option (**b.**) and attribute (**c.**) updating clusters, by fixation number and attribute weight. **d-e.** Modulation of the effect of sampled value on the probability to choose the currently fixated option by quantile-binned option updating (**d.**) and attribute (**e.**) updating effects. Insets show the fitted slopes (log-odds) for each quantile. Error bars represent the within-subject standard errors computed with the Cousineau-Morey correction. In accordance with MASC, only option-level updates were dynamically modulated by attribute weights with greater effects at early fixations, which decreased over time.

While MASC predicts that attribute weighting influences option-level beliefs, the hierarchical structure of the model implies that weighting operates only at this higher level and therefore does not affect the magnitude of lower-level attribute updates. Accordingly, we expected attribute weighting not to modulate neural activity associated with attribute updating. Thus, we repeated the same analysis on the time points and channels associated with attribute-level updating effects. This analysis revealed only a significant effect of Fixation Number (*F*(5,280) = 2.933, *p* = .013, η_p_^2^ = .050), which showed that peak amplitudes generally increased from the first to the last fixation (Fig. 6c). However, in line with our predictions, there was neither a main effect of Attribute Weight (*F*(2,112) = 2.527, *p* = .084, η_p_^2^ = .043), nor an interaction (*F*(10,560) = 1.222, *p* = .273, η_p_^2^ = .021, Fig. 6b).

### Only option-level updates predict subsequent choice

Having identified the dissociable signatures of attribute- and option-level belief updates, we next tested MASC’s prediction that sampled values influence final choice via their integration into option-level beliefs. Specifically, MASC predicts that the relationship between option-level updating and subsequent choice should depend on the value of the currently sampled attribute. When this sample provides positive evidence in the associated option’s favor, then larger option-level updates should be predictive of choosing the option. To test this, we extracted fixation-level amplitudes from the clusters associated with attribute- and option-level updating effects and fitted a logistic regression predicting whether the currently fixated option was ultimately chosen. The model included the attribute- and option-updating effects and the value of the currently fixated attribute, as well as interactions between the attribute value and each updating signal. As expected, sampled attribute value strongly predicted the subsequent choice (mean *β* = 0.472, *t*(56) = 35.680, *p* < .001, *d* = 4.726). Critically, we also observed a significant positive interaction between sampled value and the option-update amplitude, such that option-level updates were more predictive of choice for fixations to positively valenced attributes (mean *β* = 0.017, *t*(56) = 2.241, *p* = .015, *d* = 0.297) (Fig. 6d). In contrast, the corresponding interaction with attribute updating was not significant (mean β = 0.006, *t*(56) = 1.159, *p* = .126, *d* = 0.153) (Fig. 6e). Although not reaching significance, a direct comparison of the two interaction effects pointed in the same direction, with a numerically stronger effect for option-than attribute-level updating (mean Δβ = 0.011, *t*(56) = 1.472, *p* = .073, *d* = 0.195).

As a complementary behavioral test, we also asked whether subjective ratings acquired prior to the choice phase were predictive of choice, over and above attribute-level differences. Since each option was rated in isolation, such a relationship would provide evidence for a representation of an option’s value which is formed independent of a direct comparison with another. For each trial, we calculated the difference in standalone ratings between the two options and tested whether this difference predicted choice, while controlling for differences on each attribute separately. The effect of rating difference was significant (mean Δβ = 0.1903, *t*(56) = 5.094, *p* < .001, *d* = 0.675), indicating that isolated option evaluations explained choice variance beyond that captured by attribute-level comparisons.

Thus, while we identified signatures of both attribute- and option level belief updates, only the latter related sampled attribute values to the option that was ultimately chosen. Together with the predictive effect of isolated option ratings, these findings provide converging evidence that option-level value representations form the comparative basis of choice.

### Lateralized signatures of preference strength and direction emerge in preparatory motor areas

As a last step in the processing hierarchy, MASC tracks the preference for the currently best option by comparing its option-level belief distribution with the competitor option to determine whether a choice is warranted or instead additional samples should be taken. Correspondingly, we defined the preference predictor as the Jeffrey’s distance between the two posterior option distributions at each fixation, signed according to the option with the higher posterior mean. Based on prior work^23,63^, we expected to see fixation-level evidence of an emerging preference, manifesting as preparatory activity over motor areas contralateral to the responding hand associated with the currently preferred option.

As expected, we found two clusters of lateralized activity, the sign of which was determined by the currently preferred option (Fig. 4d). Specifically, negative-going deflections reminiscent of the lateralized readiness potential (LRP)^64^ were observed in the hemisphere contralateral to the currently preferred option. Importantly, this signature of emerging preference was distinct from the eventual response itself, as our analysis excluded fixations that overlapped with the response, and because response onsets were modelled as separate events in the same deconvolution model (see Methods).

In a next step, we added the final Choice as an additional regressor alongside the Preference predictor in the regression model. This analysis served two purposes. First, it can provide further evidence for a successful separation of neural signatures of the emerging preference from those of the eventual choice. Second, it allows distinguishing effects related to the direction of evidence (i.e., left vs. right) from the magnitude of preference (as for the majority of fixations, the currently best option was also the eventually chosen option). Although attenuated, the significant effects of Preference persisted when including Choice in the model. Interestingly, evidence of lateralization relating to the Choice predictor emerged relatively early and remained sustained across the time window (Fig. 7a), whereas residual Preference effects were confined to later time points (Fig. 7b). This might suggest that the direction of preference (i.e., left vs. right) is maintained over consecutive fixations, while the strength of this preference is directly modulated by the momentary changes after encoding the attribute- and option-level belief updates by the current fixation (see above).

**Fig. 7.**
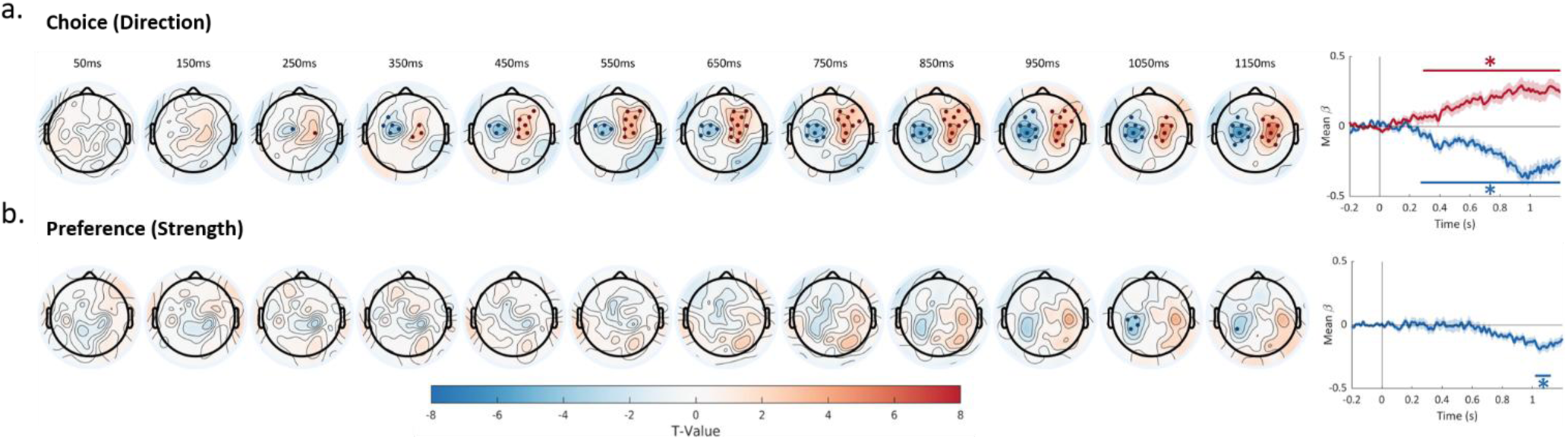
Separating the direction and strength of the emerging preference. **a.** The direction of current preference, accounted for by the Choice predictor is reflected in lateralized and sustained activity across the fixation window. **b.** In contrast, the magnitude of the preference, reflected in the residual variance accounted for by the Preference predictor appears comparatively late, presumably after the belief update provided by the current fixation is encoded.

## Discussion

Despite the ubiquitous challenge to consider many sources of information in everyday human decision-making, the neural processes underlying these choices remain elusive. A fundamental question is whether the brain integrates information to compute and compare the value of options, a core assumption in decision neuroscience that has recently been challenged both conceptually ^18,19,22^ and empirically ^31,35^. Here, we provide behavioral, computational and neural evidence that multi-attribute decisions are governed by hierarchical belief updating processes in which attribute-level beliefs are integrated into option-level representations that form the comparative basis of choice. To study these processes, we combined EEG and eye-tracking to isolate fixation-evoked neural activity in a choice paradigm that allowed participants to freely move their eyes and to respond at will. Taken together, our work sheds light on the question of neural value computation and establishes a framework for future research on the neural basis of attention and choice under naturalistic viewing conditions.

At the core of our computational framework (MASC^21^) is the assumption that multi-attribute decisions unfold through an iterative, hierarchical process, in which beliefs about the value of individual attributes are updated with each new information sample, before being weighted and integrated into option-level value representations. Consistent with this architecture, we found dissociable effects of attribute-level and option-level updating with distinct spatial profiles. Attribute updates were reflected in a positive fronto-central response that scaled linearly with update magnitude, while the integration of these beliefs into option-level representations was associated with a positive cluster over centro-parietal electrodes. Furthermore, option-level, but not attribute-level, update effects were sensitive to the weight of the fixated attribute, with fixations to highly-weighted attributes producing larger option-update responses during early fixations, before this effect decayed over time, as predicted by MASC. Critically, the functional relevance of this distinction was also evident in participants’ subsequent choices. As predicted by MASC, option-level updating effects interacted with the value of the currently sampled attribute in predicting whether the currently fixation option was ultimately chosen. This pattern is consistent with MASC’s proposed hierarchy, in which sampled values first update attribute-level beliefs, but exert their influence on choice through their integration into option-level value representations. Notably, our findings of spatially dissociable neural signals of attribute and option-level updating are consistent with fMRI research, which has established a comparable separated encoding of attribute and integrated option values in distinct brain regions^30^.

MASC’s architecture of fixation-dependent hierarchical belief-updating processes distinguishes it from many other theories of multi-attribute decision-making. For instance, the multi-attribute attentional drift-diffusion model assumes more gradual preference-formation dynamics, in which new fixations only bias the drift of an otherwise continuous evidence trace^10^. Most notably, MASC is distinct from accounts that assume multi-attribute decisions to emerge without any computation of integrated option values^18,19^. These accounts are supported by behavioral and neural evidence that multi-attribute decisions can rely on direct attribute-level comparisons^2,31,33^. Most directly, a recent study in non-human primates found that orbitofrontal neurons encode gaze-dependent comparisons between attributes rather than integrated option values^35^ (but see ^26,27^). These divergent findings can be reconciled by noting that a hierarchical architecture at the level of the whole decision is not necessarily incompatible with attribute-wise comparison processes in specific regions or cell populations that feed into it. In its current form, MASC does not directly implement such attribute-level comparison processes, although the model does account for the attentional prioritization of the most important attribute at early fixations (Fig. 3e) and the associated FEP patterns (Fig. 6b). Importantly, participants in this study and our previous work^21^ exhibited an overall tendency towards option-wise search, suggesting that decisions were mainly based on value integration (although attributes of the same option were spatially closer together in the present study to allow measuring lateralized preference signals, we obtained highly similar search patterns in our previous work^21^, which used a distance-controlled design). Yet, future extensions of our computational framework could enable attribute-wise comparisons to test how they can impact choices, possibly through influencing the neural computation of option values.

Notably, the FEPs representing attribute and option-level belief updates in the present study resemble previously characterized ERPs, situating them within existing frameworks. For attribute updates, the spatiotemporal profile of the effect is reminiscent of the P3 event-related potential^58^, which is commonly divided into an earlier fronto-central (P3a) and a later parietal (P3b) component. Given the frontal topography of the effect and its timing, the present findings appear more compatible with P3a. P3a amplitude has been shown to scale with the magnitude of belief updates during Bayesian inference and learning^59,60^, while work using fMRI has shown that model-derived measures of Bayesian belief updating are correlated with activity in the anterior cingulate cortex^65^, an area proposed to be the cortical generator of the P3a^66^. In contrast, the effect associated with option updating is highly similar to the centro-parietal positivity (CPP)^67^, a component strongly implicated in both perceptual and value-based decision-making^24,28,68–70^. While the CPP has traditionally been linked to evidence accumulation, acting as an integrator over previous decision samples^24,61,62,71^, other work using tasks in which information is sampled sequentially has suggested that the CPP indexes belief updating. For example, in a perceptual categorization task^72^, a CPP-like response observed around 500 ms after sample onset was found to encode ‘decision updates’, showing positive encoding of the current sample alongside negative encoding of neighboring samples. Remarkably consistent with our results, a recent study^63^ used an expanded judgment task and found the CPP to reflect the magnitude of sample-level belief updates, while decision evidence was accumulated downstream in preparatory motor areas. Accordingly, it was proposed that the CPP reflects the intermediate step of mapping individual samples into decision-relevant evidence prior to its accumulation elsewhere. Supporting this interpretation, the CPP in the present study appears to be integrating attribute-level beliefs into option-level representations, whereas later activity in preparatory motor areas tracks the MASC-generated Preference predictor, which reflects the evolving evidence in favor of an option and the currently preferred option over successive samples.

This lateralized signature of emerging preference is consistent with earlier work linking classic EEG motor potentials to the gradual formation of value-based decisions^23,25^. It also converges with recent evidence from single-unit recording in non-human primates, where an evidence accumulation signal in the pre-supplementary motor area was found to continuously depend on the value of both the currently attended and the unattended options^73^. Together with our own finding that the preference signal reflects the evolving comparison between both options’ integrated value estimates, these convergent results across species and methodologies suggest that the motor system tracks the relative evidence between options rather than a serial accept/reject architecture in which choice arises from evaluating one option at a time^74^.

Although the fixated value predictor is not specific to MASC, the representation of sampled values can be considered its first processing step. Interestingly, we found the associated transient and fronto-central potential to be disproportionately driven by high-value attributes, which were also fixated longer by participants. We conjecture that high-value attributes are preferentially processed, as they allow orienting future attention to candidate options that warrant further inspection (as predicted by MASC). Preferential processing of high-value attributes is also consistent with work on the reward positivity (RewP), an ERP component commonly observed in reinforcement-learning tasks in response to positive feedback^75^. The RewP shows a similar fronto-central topography to the fixated-value effect observed here, it correlates with hedonic appraisal^76^, and it responds disproportionately to positive as compared to neutral and negative outcomes^75^. Taken together, the RewP-like fixated-value effect may reflect an early evaluative signal which indicates that the current information is promising, before this information is incorporated into a belief update.

To the best of our knowledge, this study is the first to combine EEG and eye-tracking in a free-viewing, value-based decision-making paradigm, extending a methodology that has so far been used predominantly to study reading and scene exploration^43,77,78^. By capturing both the temporal dynamics of neural processing and the real-time allocation of visual attention, the two complementary measures offer a powerful framework for investigating how information is sampled and transformed into choices under naturalistic viewing conditions. Beyond the multi-attribute decisions examined here, the same approach is broadly applicable to other choice contexts in which eye movements play a critical role, including decisions involving many alternatives^79^ or options with perceptually challenging or noisy evidence^80^, as well as multi-cue inferences and categorization^41,81^. The MASC framework may also generalize to covert attentional shifts^82^, provided these can be accurately measured. More generally, our findings establish free-viewing EEG–eye-tracking as a promising tool for probing the interaction between attention and choice, and for studying the neural computations that support a wide range of complex decisions.

In summary, by combining EEG and eye-tracking paired with the computational framework of the MASC model, we provide a unified account of the neural mechanisms underlying multi-attribute decision-making, tracing the process from information sampling to preference formation. We show that core predictions of MASC are reflected both in patterns of overt attention and in latent neural activity, which can be hierarchically dissociated into attribute and integrated option-level representations. We further show that these belief states are sequentially updated to shape preferences over successive samples. Together, our results provide evidence that multi-attribute decision-making unfolds as a recurrent and hierarchical process of sequential sampling, integration and updating.

## Methods

### Preregistration

Prior to data collection, the project was preregistered on the Open Science Framework (OSF, https://osf.io/bv6s5/), using the OSF preregistration template. We preregistered two sets of hypotheses relating to continuous signals which track the emergence of a choice throughout the trial, and markers of updating at the level of individual fixations. More specifically, we predicted to identify updating signals at the level of attributes and options, larger (option-level) updates for more important attributes, and an association of lateralized preparatory motor activity with the emerging preference. Preregistration also included exclusion criteria (i.e., incomplete datasets, poor eye-tracking and/or EEG quality) as well as the pre-processing of the EEG data and the subsequent deconvolution analysis.

### Participants

To determine the sample size, an a-priori power analysis was performed based on conducting separate tests for detecting EEG signals related to updating processes (e.g., option-level updating) and to continuous processes (e.g., emerging preferences). To achieve a joint statistical power of 80% for detecting both effects, the power analysis suggested testing at least 52 participants. Taking dropouts into account, a total of 70 participants were recruited via the University of Hamburg’s online recruiting system in exchange for either course credit or monetary remuneration (15 Euro per hour). Participants were required to be between 18 and 35 years of age, be able to speak German, not wear glasses, or suffer from any physical, psychological or neurological disorders. Data from 13 subjects was excluded due to technical issues (6 participants), excessive movement leading to poor quality EEG data (5 participants) and participants pressing buttons at random without fixating the attributes (2 participants). This left 57 participants in the final analysis (44 females, 13 males, mean age = 24.75 years, SD = 4.89 years). All participants gave written informed consent and the study was approved by the Local Ethics Committee of the Faculty of Psychology and Human Movement Sciences at the University of Hamburg (#AZ 2022_046).

### Stimuli

The study was conducted in a darkened room and stimuli were presented using Psychtoolbox version 3.0.19 in MATLAB R2021b. The background was gray (RBG: 127, 127, 127) throughout the experiment and stimuli were custom made icons (100 x 175 pixels) representing the three attributes (battery capacity, storage space and screen size), with their color denoting the option to which they belonged (green (RGB: 37, 86, 24), or blue (RGB: 0, 67, 164) in the choice task and black (RGB: 0, 0, 0) in the rating task). A pale-yellow star (RGB, 255, 255, 192, alpha = 0.76) spanning 100 x 100 pixels was superimposed onto the bottom of each attribute icon, which displayed the respective star rating in black text to one decimal place. Stimuli were presented on a 24-inch screen with a resolution of 1920 x 1080 pixels and a refresh rate of 144 Hz. Button presses were made on a mechanical keyboard with <1 ms latency.

### Procedure

The main experiment consisted of two parts, a rating phase followed by a choice phase. At the beginning of the rating phase, participants were informed about the three smartphone attributes (screen size, battery capacity and storage space), the range of star ratings (0-5) and shown the icons associated with each attribute. In each rating trial, the three attribute icons (displayed in black) together with the corresponding star ratings were displayed for a single smartphone. The attribute icons were arranged horizontally (see Fig. 2a and Supplementary Fig. 1), with their position randomized across trials. Participants were instructed to rate how much they liked the smartphone based on the displayed attributes using a mouse-controlled slider ranging from 1 to 10. As the mouse was moved along the scale, the current rating was updated and displayed in real time. Participants selected their rating by clicking the mouse, after which the selected value was shown to 1 decimal place. They could then either confirm the rating by pressing the space bar or revise it by pressing the ‘q’ key. Pressing the space bar advanced to the next trial, whereas pressing the ‘q’ key reset the current one. All participants rated the same set of 45 simulated smartphones, presented in a randomized order across participants. The attribute values of the smartphones were sampled pseudo-randomly, ensuring an equal distribution of low-value (overall value = 0-5 stars), medium-value (overall value = 5-10 stars) and high-value (overall value = 10-15) options. Neither eye movements nor EEG were recorded during the rating phase.

After completion of the rating task, a custom-built algorithm was used to construct equally sized sets of easy, medium and hard trials based on the participant’s ratings for each smartphone. First, all possible pairs of options were generated and ‘no-brainer’ trials in which one option was superior on all attributes were discarded. For the remaining valid comparisons, trials were ranked and binned into tertiles (easy, medium, hard) based on the rating differences for the option pairs. To ensure the most even distribution of option appearances, a permutation test was conducted for each difficulty level. Over 50,000 permutations, 140 comparisons were drawn at random from each difficulty bin and the standard deviation of the chosen option counts were calculated. The draw with the most even distribution of item appearances (the lowest standard deviation) was retained. This resulted in a total of 420 choice trials (140 per difficulty) and ensured each option was roughly equally represented.

In the choice phase, participants were reminded of the attribute icons, the range of star ratings and were informed that they would now choose between two smartphones in each trial (a green option and a blue option). Each trial began with a black fixation cross displayed in the center of the screen, on which participants were required to maintain fixation on for 500 ms before a trial would start. A square region of interest (400 x 400 pixels) invisible to the participant, was centered on the fixation cross. Once the eye tracker recorded that participants had maintained fixation within this region for at least 500 ms, the trial would begin. If participants moved their eyes out of this region within 500 ms, the timer was reset. This ensured that initial fixations were always in the center of the display and that the trial could not begin if the position of the eye could not be accurately registered.

Following the offset of the fixation cross, the attribute icons, without the associated star ratings appeared. The six stimuli (three attributes for each option) were positioned at equal angular intervals on an invisible circle with a radius of 400 pixels, forming a hexagonal configuration. Each attribute icon was colored based on its corresponding option (green (RGB: 37, 86, 24), or blue (RGB: 0, 67, 164)), with all attributes for the blue option presented on one side of the screen and all attributes for the green option on the other side. This provided an intuitive mapping of each option to a responding hand, which later allowed us to measure lateralized readiness potentials tracking the emerging preference for an option. The side associated with each option color was counterbalanced across participants. This preview period was intended to give participants time to identify the locations of each attribute, thereby minimizing the occurrence of early exploratory fixations related to display familiarization rather than decision processes. To further help participants localize the attributes, the positions of the attribute icons stayed constant for 70 consecutive trials, after which they were reassigned. Before each reassignment, participants were notified that the positions were to change through an onscreen message.

After 1500 ms, the star ratings appeared, superimposed onto the attribute icons. Participants were instructed to choose which option they preferred using the ‘S’ and ‘K’ keys for the left or right option using their left and right index fingers, respectively. The stimuli remained on screen until a choice was made and participants could move their eyes freely. Once a choice was made, the respective response square was highlighted with an arrow for 500 ms to inform the participant that their response had been registered, after which the screen turned blank for an intertrial interval sampled from between 1500 and 2500 ms before the presentation of the fixation cross for the next trial. Participants were invited to take a break every 50 trials. If a participant decided to take a break, they were required to press the ‘S’ key to indicate when they were ready to continue and the eye tracker was recalibrated.

### Statistical analyses

For the behavioral analysis of choice and attention patterns, a frequentist statistical approach was used. Data was aggregated within participants (e.g., means and regression coefficients) and inferential statistics (e.g., *t*-tests, Pearson correlation) were performed at the group level using an alpha-level of 5%. All behavioral *t*-tests were one-sided. For the threshold-free cluster permutation tests, significant clusters were identified using a two-sided (corrected) significance threshold of *p <* .05. Contiguous significant samples were grouped into clusters based on the predefined electrode neighborhood structure.

### Model Fitting

The MASC model assumes that multi-attribute decisions unfold through an iterative process of sampling and updating beliefs about the value of attributes and options. At each step, fixation-derived information samples are used to update attribute-level belief distributions, which are weighted by attribute importance and integrated to form higher-level option belief distributions. This Bayesian framework allows examining how a decision develops over time and generating predictions about the magnitude of belief updates by tracking how these distributions evolve with each successive sample.

To derive the model-generated estimates used to predict fixation-evoked activity, we first fit the MASC model to the behavioral data. To determine attribute weights for each participant, we used a general linear model to predict the participant’s option ratings from the normalized values of each attribute. The beta weights for each participant were normalized to ensure they summed to 1, a prerequisite for the model. In cases where the beta weights were negative, the values for the corresponding attribute were multiplied by -1. We fit the model to each participant to obtain the best-fitting estimates for the model’s three free parameters i.) standard deviation of sampling *σ_s_*, ii.) increase (per fixation) of tolerance Δ, and iii.) sensitivity of search rule α while keeping the initial tolerance θ_0_ fixed to 0.1. The best-fitting estimates were obtained using a grid search with 31 equidistant values for each parameter (*σ_s_*: from 0.001 to 3.001; Δ: from .01 to .09; α from 0 to 5) in which model fit was approximated with a set of summary statistics: the proportion of trials in which the model correctly predicted the consistency with which participants chose the higher value option, the difference between the observed and predicted proportion of fixations to each option-attribute pair, and the difference between the observed and predicted number of fixations per trial. Note that the latter summary statistic was normalized by the maximum observed number of fixations for each participant to constrain the statistic to values between 0 and 1. We identified the best-fitting parameter set as the set with the highest mean score across these summary statistics.

### Eye-movement recording

Eye movements were recorded monocularly using a video-based EyeLink 1000 Plus eye tracker (SR Research Ltd.) with a sampling rate of 1,000 Hz. Head position was stabilized using a chin and forehead rest positioned at a distance of 92 cm from the screen with the eye tracker in between. The eye tracker was calibrated at the beginning of the experiment using a 9-point calibration layout to ensure a maximum calibration error of < 1.0° and recalibrated regularly throughout. For all participants, the right eye was tracked.

### EEG data acquisition

Electrophysiological data were recorded from 64 active electrodes using Brain Vision Recorder (Brain Products) and referenced online against Cz at a sampling rate of 1000 Hz. 63 electrodes were mounted in a custom layout (see Supplementary Fig. 2) on a textile cap (ActiCap, Brain Products), while the additional electrode was placed infraorbitally beneath the right eye to record EOG activity. Impedances were kept below 20 kΩ.

### EEG and eye-movement preprocessing

Preprocessing of the EEG data was performed in MATLAB R2024b (The Mathworks Inc. 2024) using the EEGLAB^83^ toolbox and the EYE-EEG^43^ plugin. A schematic diagram of the preprocessing and analysis pipeline is shown in Fig. 2. Both the EEG and eye tracker were originally recorded at a sampling rate of 1000 Hz but were downsampled offline to 500 Hz. During the choice phase, stimulus and response onset triggers were sent simultaneously to both the eye tracker and EEG system, as well as triggers marking the start and end of the phase (Fig. 2b). Using the EYE-EEG toolbox, these triggers were used to synchronize events across the EEG and eye-tracking data. The mean synchronization error across all participants and events was < 2 ms.

Next, saccade and fixation events were identified and imported using the Engbert & Kliegl (2003)^84^ algorithm for saccade detection as implemented in the EYE-EEG toolbox. Saccades were identified using a velocity threshold of 5 median based standard deviations and were required to have a minimum duration of 5 ms. Saccades occurring within 25 ms of one another were merged, and only the first saccade of the group was retained. Fixation onset and offset were defined by the offset of the preceding saccade and the onset of the subsequent saccade, respectively. Time periods in which gaze fell outside the screen boundaries (horizontal position > 1920 or vertical position > 1080) were flagged and excluded from all subsequent analyses. To determine areas of interest (AOI) we defined a circular region (radius of 200 pixels) around each attribute location. Any fixations falling within this region were recorded as fixations to the corresponding attribute, while fixations which could not be assigned to any AOI were discarded.

The downsampled EEG data was initially high pass filtered at 0.1 Hz with a second-order Butterworth filter and subsequently re-referenced to average. Noisy channels were detected using a voltage threshold of five standard deviations and then spherically interpolated. To remove ocular artefacts, we employed an optimized ICA algorithm (OPTICAT^46^), an approach which addresses several limitations of standard ICA when used with free-viewing data. In particular, standard ICA often fails to identify the saccadic spike potential (SP), a brief, high-frequency electromyographic signal generated by the extraocular muscles at saccade onset^46^. Due to its short duration (∼20 ms), varying topography depending on saccade direction and relatively small amplitude compared to other ocular artefacts, it is challenging for conventional ICA to reliably detect and remove it. To account for the SP’s overall weak contribution to the signal, the OPTICAT approach improves detection by overweighting SPs in the training data. Following the recommendations provided by Dimigen (2020)^46^, the training data was band-pass filtered with edges 2 Hz-100 Hz before 30 ms intervals ([-20 ms, +10 ms]) were taken around each saccade onset and repeatedly appended to the EEG data until the data was twice its original length. The right EOG channel was included in the training data, and periods with extreme movement artefacts (>500 uV) were removed. Finally, artefactual components were flagged and later removed using an eye-tracker guided artefact identification algorithm^36^ with a variance ratio threshold of 1.1. We also used the ICLabel^85^ plugin to classify components and removed any remaining components that were determined as eye-related with more than 90% confidence.

### Linear deconvolution modeling

Beyond the correction of ocular artefacts, two major challenges remain when studying FEPs. First, fixations occur in rapid succession, typically 3-4 times per second, meaning that the neural activity elicited by each fixation temporally overlaps. In turn, the traditional ERP approach of extracting a window time-locked to fixation onset does not cleanly isolate activity attributable to one event. Secondly, FEPs are sensitive to factors such as fixation duration and saccade amplitude^47,51–54^. If these features are not appropriately controlled for, they can produce spurious effects that appear cognitive in nature. However, advances in the analysis of combined EEG and eye-tracking have allowed these issues to be addressed simultaneously through the use of linear deconvolution modeling^44,47,49^. By assuming that the observed EEG activity is a linear summation of activity from neighboring events, the signal can be deconvolved to isolate non-overlapping activity associated with the current event of interest (i.e., single fixations). Moreover, the dependence of fixation-evoked activity on fixation duration and saccade amplitude can be addressed by including these predictors as covariates in the model. Here we used deconvolution modeling as implemented in the Unfold.jl (v0.8.9) toolbox^44^ to model fixation-evoked activity as a function of MASC-generated predictors, while simultaneously controlling for temporal overlap and the non-linear effect of saccade amplitude and fixation duration.

To derive the MASC-generated predictors, the model was simulated with the best fitting parameters. For each participant, trial, and fixation, we provided MASC with the observed fixated attribute and updated the model’s belief distributions accordingly. This allowed us to obtain an estimate of the predicted strength of belief-updating at each fixation, given the parameterization and the participant’s fixation path. Within MASC, updating a belief about the value of an item is formalized as a shift in the (normally-distributed) posterior belief distribution and is observable at both the level of individual attributes (*AttUpd*) and of options (*OptUpd*). Therefore, to quantify ‘belief-updating’ at each level, we calculated the MASC-predicted KL-divergence between the posterior belief distribution pre (*X_t_*_−1_) and post (*X_t_*) fixation as follows.

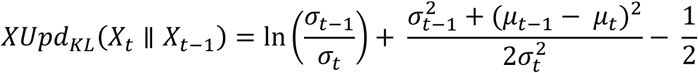

where *X* refers to the fixated attribute posterior in the attribute-level case and the fixated option posterior in the option-level case. Since KL divergence is sensitive to differences in both location and scale, this allowed us to jointly quantify not only changes in the belief about the average value of the fixated item, but also the corresponding change in uncertainty.

We further hypothesized that the distance between the two option posterior distributions would modulate the FEPs. Prior work has demonstrated that preparatory motor signals reflecting a developing propensity to respond emerge not only at the time of the final decision, but already during the processing of individual information samples^23^. Within the framework of MASC, the separation between option posteriors can be interpreted as a propensity to respond given the sampled evidence, as the stopping rule specifies that a choice is triggered once the overlap between option posteriors falls below the tolerance parameter θ. Accordingly, we also calculated a third *‘Preference’* measure as the Jeffrey’s distance (i.e., the sum of asymmetric KL divergences) between the option posterior distributions at each fixation and coded it according to the option distribution with the currently highest posterior mean.

Finally, while our main focus was fixation-evoked potentials, we also included two terms in the same model to account for the influence of stimulus onsets and the button-press response. Namely, we modelled stimulus-evoked potentials as intercepts to account for their overlap with early fixations and responses as a function of responding hand.

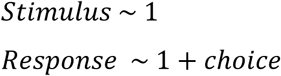

To further separate fixation-evoked activity from response-activity, any fixations which temporally overlapped with the button-press response were removed from analysis (5.4% of all fixations). The deconvolution models were fit to the continuous, pre-processed EEG data of each participant separately, before the resulting betas were subject to threshold-free cluster permutation tests (*E* = 0.5, *H* = 2, N = 5000) implemented using the ept_TFCE toolbox^56^. Clusters were identified using the calculateClusters.m function with a threshold of .001. In the design matrix, stimulus and fixation onsets were time-expanded over a window of -200 to 1,200 ms, while response onsets were time-expanded over a window of -1000 to 200 ms.

## Data availability

All behavioral data is publicly available at https://github.com/jordandeakin/MASC-EEG

## Code availability

The analysis code to produce all main results and figures is publicly available at https://github.com/jordandeakin/MASC-EEG.

## Acknowledgements/Funding

We thank Izel Habil, Changfa Fu and Amir Hossein Hadian Rasanan for help with data collection and the Cognitive Modelling & Decision Neuroscience team at the University of Hamburg for many insightful discussions. This work was supported by a collaborative research grant, co-funded by the German Research Foundation / Deutsche Forschungsgemeinschaft (DFG; grant no. GL 984/1-1) and a collaborative research grant, co-funded by the German Ministry of Education and Research / Bundesministerium für Bildung und Forschung (BMBF; grant no. 01GQ2303). S.G. also acknowledges support by the European Research Council (ERC) under the European Union’s Horizon 2020 research and innovation program (Grant agreement No. 948545.

## Author contributions

J.D. and S.G. conceptualized the study and developed the theoretical framework. J.D. designed and programmed the task and collected and analyzed the data. R.F. contributed to EEG methodology, task design, data analysis, and interpretation of the results. J.D. wrote the original draft of the manuscript. S.G. and R.F. reviewed and edited the manuscript. All authors contributed conceptual input. S.G. supervised the project and acquired funding.

## Competing Interests

All authors declare no competing interests.

## Notes

https://github.com/jordandeakin/MASC-EEG

### Competing Interest Statement

The authors have declared no competing interest.

### Summary of Updates

Additional analyses and figures have been added to explore how the identified attribute- and option-level update signals predict choice behaviour.

