## Supplementary Material for "Fixation-evoked potentials reveal neural signatures of hierarchical value-integration during decision-making"

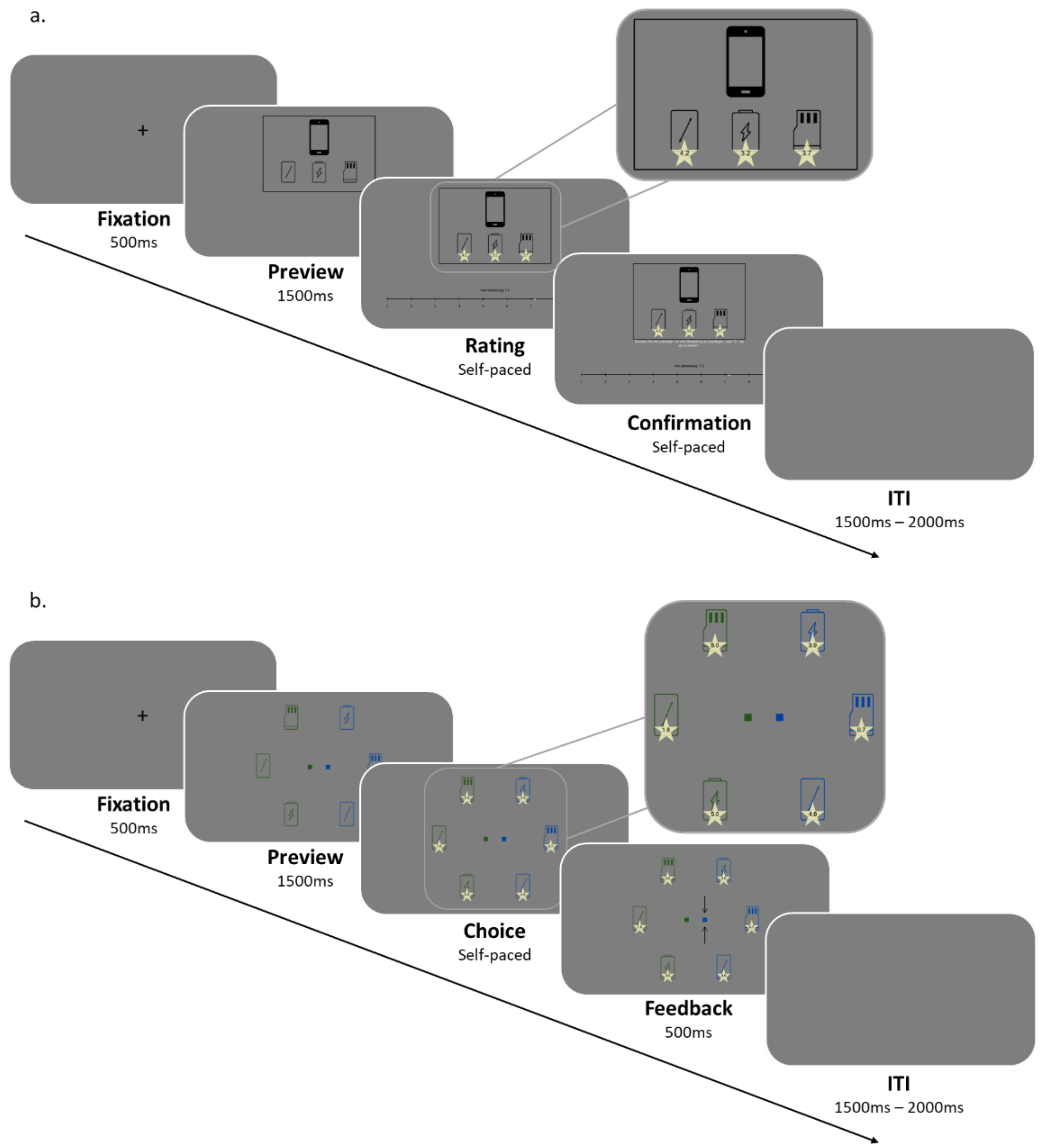


### Fig S1 | Trial structure of rating and choice phases

**a.** Trial structure for the rating task. *Fixation:* A fixation cross was presented for 500 ms. *Preview:* Attribute icons were shown without star ratings for 1500 ms. *Rating:* Star ratings and a 1–10 rating scale appeared. Participants rated the smartphone using the mouse-controlled slider. *Confirmation:* Participants confirmed their response with the space key or could revised it with the 'q' key. *Intertrial interval (ITI):* A blank screen was presented for 1500–2000 ms before the next trial. **b.** Trial structure for the choice task. *Fixation:* A fixation cross was presented for 500 ms. *Preview:* Six attribute icons (three per option) were shown without ratings for 1500 ms. *Choice:* Participants selected their preferred option using the 's' (left) or 'k' (right) key. *Feedback:* An arrow indicated the chosen option. *Intertrial interval (ITI):* A blank screen was presented for 1500–2000 ms before the next trial.

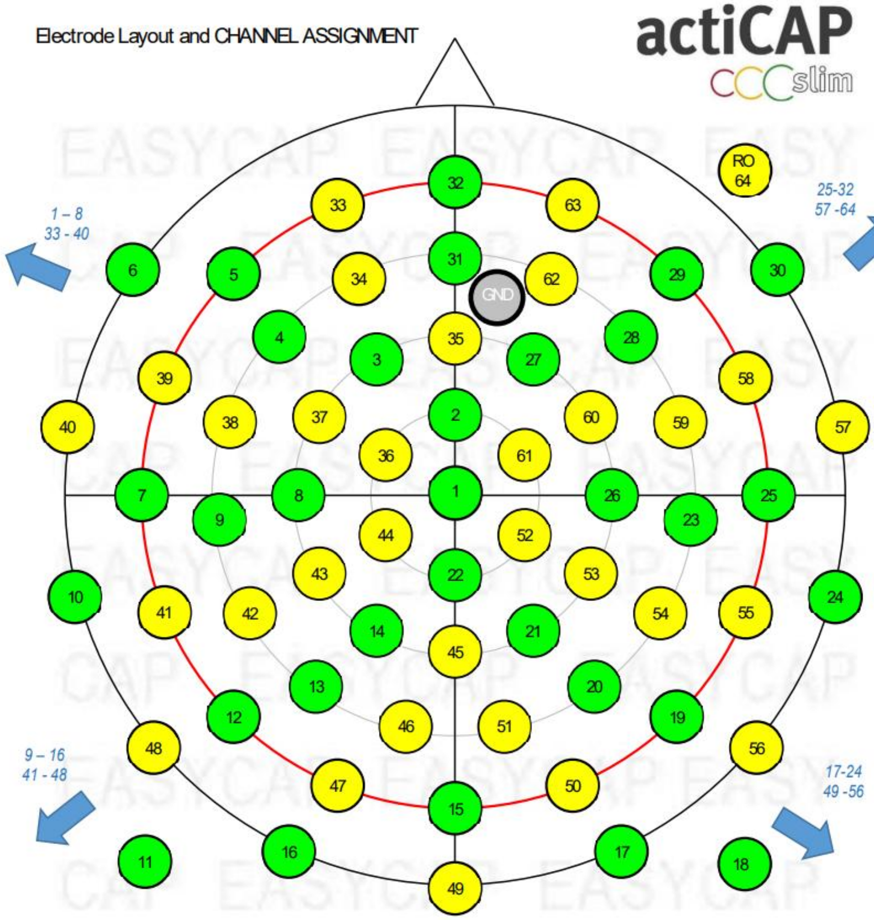


**Fig. S2 | Custom ActiCap (Brain Products) layout**

The montage places 61 electrode positions onto 5 equidistantly spaced concentric rings around Cz (1). The positions on each ring are also equidistantly from each other. The positions on the vertical and horizontal central lines are identical to 10%- positions. The ring marked red is defined as Theta = 90°and passes through Fpz (32), T8 (25), Oz (15), T7 (7). The outmost ring reaches from the canthii to the Inion

| Predictor | Sum of Squares (Type III) | *df* | Mean Square | *F* | *p* | partial $\eta^2$ |
|---|---|---|---|---|---|---|
| Fixated Value | 494.107 | 3.385 | 128.834 | 11.085 | < .001*** | 0.165 |
| Error | 2,496.171 | 214.772 | 11.622 | | | |

**Table. S1 | One-Way Analysis of Variance for the effect of Fixated Value on fixation duration.**
Degrees of freedom were corrected using Greenhouse-Geisser estimates of sphericity (ε = 0.77)

| Comparison (Rounded Value, V) | Estimate | SE | *df* | *t* | *p* | | Cohen's d |
|---|---|---|---|---|---|---|---|
| V1 – V0 | -0.481 | 0.570 | 56 | -0.845 | .402 | | -0.038 |
| V2 – V0, V1 | 0.045 | 0.492 | 56 | 0.092 | .927 | | 0.004 |
| V3 – V0, V1, V2 | 1.012 | 0.409 | 56 | 2.473 | .016 | * | 0.081 |
| V4 – V0, V1, V2, V3 | 2.106 | 0.408 | 56 | 5.159 | < .001 | *** | 0.168 |
| V5 – V0, V1, V2, V3, V4 | 2.254 | 0.489 | 56 | 4.607 | < .001 | *** | 0.180 |

**Table. S2 | Contrasts for the effect of Fixated Value on fixation duration.**
Rounded Value refers to the rounded star rating of the fixated attribute (e.g., V3 = 3 stars).
